# Packpred: Predicting the functional effect of missense mutations

**DOI:** 10.1101/2020.12.30.424909

**Authors:** Kuan Pern Tan, Tejashree Rajaram Kanitkar, Kwoh Chee Keong, M.S. Madhusudhan

## Abstract

Predicting the functional consequences of single point mutations has relevance to protein function annotation and to clinical analysis/diagnosis. We developed and tested Packpred that makes use of a multi-body clique statistical potential in combination with a depth dependent amino acid substitution matrix (FADHM) and positional Shannon Entropy to predict the functional consequences of point mutations in proteins. Parameters were trained over a saturation mutagenesis data set of T4-lysozyme (1966 mutations). The method was tested over another saturation mutagenesis data set (CcdB; 1534 mutations) and the Missense3D data set (4099 mutations). The performance of Packpred was compared against those of six other contemporary methods. With MCC values of 0.42, 0.47 and 0.36 on the training and testing data sets respectively, Packpred outperforms all method in all data sets, with the exception of marginally underperforming to FADHM in the CcdB data set. On analyzing the results, we could build meta servers that chose best performing methods of wild type amino acids and for wild type-mutant amino acid pairs. This lead to an increase of MCC value of 0.40 and 0.51 for the two meta predictors respectively on the Missense3D data set. We conjecture that it is possible to improve accuracy with better meta predictors as among the 7 methods compared, at the least one method or another is able to correctly predict ∼99% of the data.

## 2. Introduction

Amino acid substitutions could affect protein stability, alter/impair its function and possibly lead to disease conditions (Zhang et al., 2012). Several such single amino acid substitutions in proteins, also called missense mutations, are implicated in diseases such as cystic fibrosis, diabetes, cancer etc. (Roach et al., 2010; Stranger et al., 2011). Data from clinical studies as well as from large-scale projects such as the Human Genome Project (Craig Venter et al., 2001), HapMap Project (Frazer et al., 2007), Exome Sequencing Project and the 1000 Genomes Project (Altshuler et al., 2012) unearth such single amino acid mutations. It would be instrumental to have a fast and automated computational method to accurately predict the functional effect of these mutations. Such an exercise could also provide valuable insights into the development of personalized medicine.

Several computational methods predict the effect of missense mutations. The methods utilize sequence or structure information or a combination of the two. The sequence based methods rely on previously known protein sequences and their characterizations deposited in databases. For example, in the SIFT method (Ng and Henikoff, 2003), mutational effect prediction are made based on a customized position specific substitution matrix (PSSM), constructed with PSI-BLAST (Altschul et al., 1997) and MOTIF finder (Smith et al., 1990) to identify conserved local sequence regions. A majority of structure-based methods are based on machine learning algorithms. These methods employ different feature sets and machine learning architectures. For example, I-mutant2.0 (Capriotti et al., 2005) is trained on features such as pH, temperature and mutation type using a support vector machine. AUTO-MUTE 2.0 (Masso and Vaisman, 2014) constructs a statistical contact potential with Delaunay tessellation and trained their models with additional attributes such as ordered identities of amino acids, pH and temperature. PoPMuSiC-2.0 (Dehouck et al., 2009) uses a linear combination of 26 different statistical energy functions in an artificial neural network architecture. mCSM (Pires et al., 2014b) utilizes a graph metric to summarize physicochemical interactions within a cut-off distance as pattern signatures, and trained them with Gaussian process regression model. SDM (Pandurangan et al., 2017), which does not rely on machine learning, constructs an environment-specific amino acid substitution matrix based on observed substitutions in evolutionary time. DUET (Pires et al., 2014a) is a meta-algorithm that consolidates the methods of mCSM and SDM (Worth et al., 2011). Missense3D (Ittisoponpisan et al., 2019) is another structure based method that uses seventeen structural properties to predict the effect of mutation. Dynamut2.0 (Rodrigues et al., 2020) uses normal mode analysis and graph-based signatures. Polyphen (Adzhubei et al., 2010), is a hybrid method that combines sequence and structural features to predict the effect of a mutation. It uses an improved version of PSSM, information from the Pfam database and structural features such as accessible surface area and volume of an amino acid to make a prediction. SuSPect (Yates et al., 2014), is another hybrid based method that uses PSSMs, Pfam domain profiles(Finn et al., 2014). It also includes information from protein-protein interaction network, and searches in the database for known functional annotations of a mutated position. Despite these various efforts and algorithms, the functional fate of point mutations remains a challenging problem.

A missense mutation could lead to functional instability by either disrupting its structure or by affecting its interaction interface and/or active sites without necessarily impacting its structure. A mutational effect predictor should hence take into account both, the effect of mutation on overall structural stability and on its functional relevance. In this study, we describe Packpred that addresses both these aspects. For structural features Packpred uses an environment-dependent multi-body statistical potential and a depth dependent substitution matrix, FADHM. We had previously established that FADHM scores are useful in predicting the effects of point mutations (Farheen et al., 2017). The multi-body statistical potential considers the observed/expected ratio of cliques of residues. The greater the value of the ratio, the more energetically stable is the packing of amino acids in the residue clique. We further categorized these residue cliques based on their residue depths. Residue depth (Chakravarty and Varadarajan, 1999; Tan et al., 2011, 2013) measures the degree of burial and hence the solvation effect on amino acids. Depth has been shown to correlate well with structural stability and free energy change of cavity-creating mutations in globular proteins (Chakravarty and Varadarajan, 1999; Tan et al., 2011). Our depth based statistical potential hence assesses the effect of mutation on local packing stability. To capture the functional relevance of amino acids, we used residue position Shannon entropy from a multiple sequence alignment of homologs of the query sequence. By this, we exploit evolutionary information to quantify degree of observed variation at the position of mutation. Usually, the lesser the variation the greater is the functional importance of the residue.

## 3. Materials and methods

### 3.1. Data sets

#### 3.1.1 Statistical potential data set

A set of 3753 protein structures obtained from the Protein data bank (PDB) (Berman et al., 2000) was used to construct the clique statistical potential. The structures in this set have a resolution of 2.5 Å or better, R-free of 0.25 or better and are non-redundant at 30% sequence identity. To account for atomic position fluctuations (protein dynamics) while considering amino acid cliques, 10 homology models were built using Modeller (Šali and Blundell, 1993) with the native protein serving as target and template. These homology models along with the native structure (i.e. 11 structures for each protein) were used to build a statistical potential.

#### 3.1.2 Saturation mutagenesis data sets

Saturation mutagenesis data sets of two proteins, T4-lysozyme (Rennell et al., 1991) and Controller of cell division or death B (CcdB) (Adkar et al., 2012) were used in this study. T4 Lysozyme is a 164 amino acid residue protein with our reference structure being PDB: 2LZM that was solved at a resolution of 1.7 Å (Weaver and Matthews, 1987). Each position except the first was mutated to 13 other amino acids (A, C, E, F, G, H, K, L, P, Q, R, S, and T). After excluding key catalytic site residues (D10, E11, R145, R148P) the data set consists of 1966 mutations. CcdB is a cytotoxin (an inhibitor of DNA gyrase) with 101 amino acids. Its native structure was solved at 1.4 Å resolution (PDB: 3VUB (Loris et al., 1999)). Full saturation mutagenesis (mutating each position to all other 19 amino acids) was performed at all positions of the protein. After removal of active site residues (I24, I25, N95, F98, W99, G100, I101), a final set of 1534 mutations was obtained. In both saturation mutagenesis experiments, an assessment was made on the phenotypic effect for each mutation. For T4-lysozyme the phenotypic effect was gauged based on the plaque-forming ability of the mutant. Subject to the same experiment condition, a mutant is assigned to one of the four levels of sensitivity if the size of the plaque is (i) similar to native control, (ii) significantly smaller, (iii) with hazy morphology or difficulty in discerning plaques and (iv) no plaque formation(Rennell et al., 1991). For CcdB, the mutational sensitivity score was quantitatively defined as the titer number at which the protein activity (in this case, inducing cell death) decreases by 5-fold or more relative to its previous dilution. Values of mutational sensitivity range from 2 to 9 in CcdB and we scaled the T4-lysozyme values to range from 2 to 5. For both data sets, a mutation is regarded as neutral if there is no perceptible phenotypic difference as compared to its native sequence (MS score = 2 in CcdB and T4-lysozyme), and regarded as destabilizing otherwise.

#### 3.1.3 Missense3D data set

The Missense3D data set consists 4099 mutations from 606 proteins extracted from Humsavar (Bateman et al., 2017), ClinVar (Landrum et al., 2014), and ExAC (Karczewski et al., 2017) (Ittisoponpisan et al., 2019). Humsavar lists all the annotated missense variants from humans reported in UniProt and SwissProtKB. ClinVar catalogs variations in humans and their associated phenotype. ExAC is exome aggregation consortium that describes the aggregation and analysis of human exome. The analysis includes quantification of the pathogenecity of variants. The data set of 4099 mutations consists of 1965 disease-associated variants and 2134 neutral variants (not associated with any known disease, yet). Packpred parameters were trained on the T4-lysozyme data set and tested on the CcdB and Missesense3D data sets.

### 3.2 Structural and Sequential features

#### 3.2.1. Residue depth

Depth is defined as the distance of a protein atom to the nearest bulk water molecule (Chakravarty and Varadarajan, 1999). The quantity measures the degree of burial of the atom. Depth has been shown capable of concisely describing protein environment, as substantiated by its utilities in protein design and function predictions (Tan et al., 2011, 2013, Farheen et al., 2017). Atom depth values were computed using default parameters. The depth of a residue clique is defined as the average depths of its constituent atoms.

#### 3.2.2. Cliques of amino acid residues

A clique is defined as a sub-graph in which all possible pairs of vertices are linked. We define a (N, d_cut_) “residue clique” to be a clique of N amino acids within a linkage distance of d_cut_. We consider two amino acids as linked when at least four, or more than half of side chain non-hydrogen atoms (whichever smaller) are within *d*_*cut*_ from atoms of another amino acid (Figure 1). For Glycine, the C^°^ atom is used in lieu of the side chain. Residue cliques defined with different combinations of N and d_cut_ (N ranges from 2 to 4, d_cut_ ranges from 7.0 Å to 10.5 Å in step of 0.5 Å) have been computed and investigated in this study.

**Figure 1:**
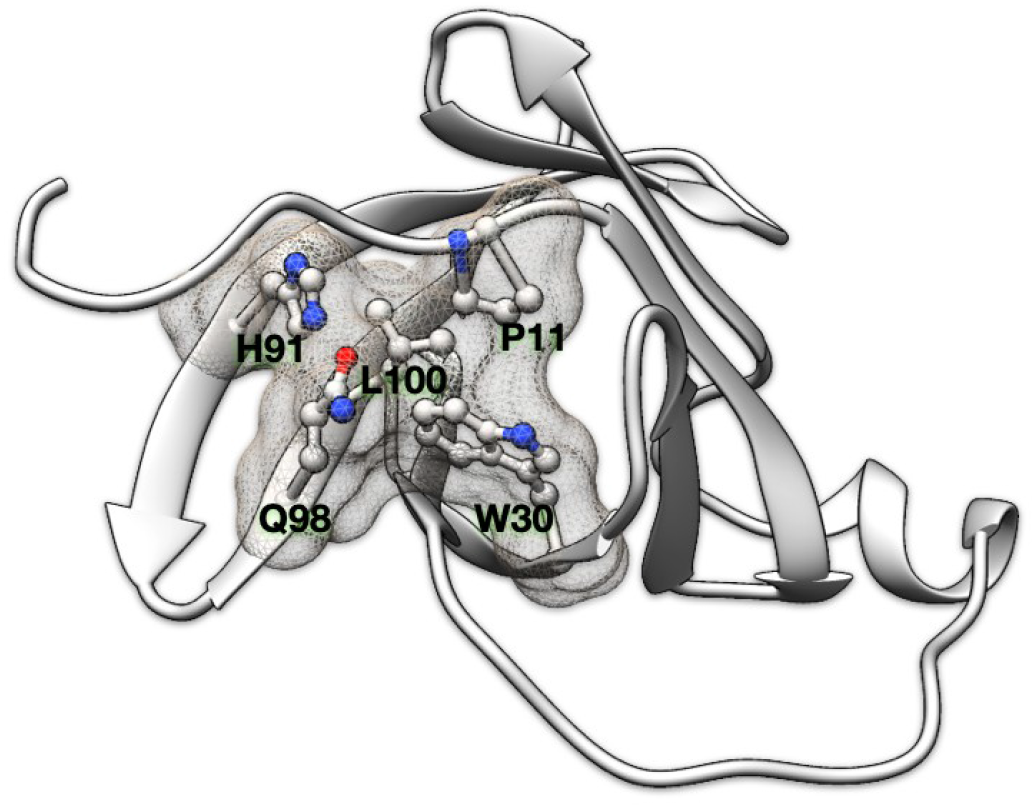
Residue clique of amino acids. (A). A 5-residue clique (P11, W30, H91, Q98, L100) of cut-off 7.5Å shown in ball and stick representation and enveloped with a meshed molecular surface from human recombinant MTCP-1 protein (PDB: 1A1X).

#### 3.2.3 Statistical potential and residue clique score

A residue clique statistical potential is constructed by adopting the formulation of Sippl’s potential of mean force (Sippl, 1990),

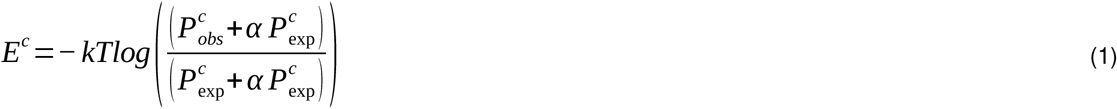

Where *E*^*c*^ is the pseudo potential energy and *c* is a residue clique of type {*r*_*1*_, *r*_*2*_, …}, where the *r*_*i*_’s are the amino acid types; P^c^ _obs_ is the observed number of residue clique c; P^c^ _exp_is its expected number in a hypothetical reference state without energetic interactions; *α* is the ratio of pseudo-count introduced to account for sparse statistics, and is taken as 0.00 in our study. -*kT* is a constant and is assumed as 1 in this study.

For each (*N, d*_*cut*_) clique, the statistical potential is built at 5 different levels of depth (2.80 Å – 5.25 Å, 4.25 Å – 6.25 Å, 5.25 Å – 7.25 Å, 6.25 Å – 8.25 Å, 7.25 Å - ∞). To calculate the score of a residue clique (S), the mean *μ* and standard deviation *σ* of its depth is first computed. A Gaussian probability density function *N*(*x* | *μ, σ*) is then accordingly built. The clique score is computed as the weighted sum of integrand at every depth level as,

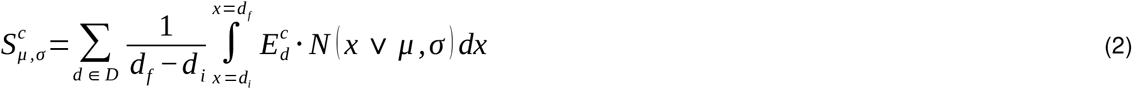

Where *d* is one depth level, *d*_*i*_ and *d*_*f*_ is the lower and upper bound of the level.

Most residue cliques in a protein are overlapping with one another, and an amino acid residue can participate in multiple cliques. The score of a residue is taken as the average of all such cliques. The score of a protein is further taken as the average of all its residue scores.

#### 3.2.4. Shannon Entropy

Shannon entropy (H) is a measure of variation observed at a given position. It calculated from a multiple sequence alignment obtained by a PSI-BLAST search against the uniref50 database (Altschul et al., 1997). H for a given position is then calculated as,

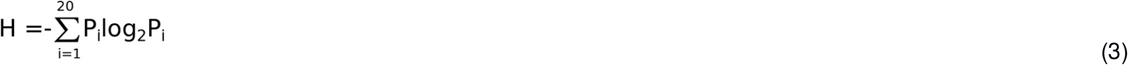

Where Pi is the fraction of amino acid i observed at a given position.

#### 3.2.5. FADHM Scores

FADHM scores are depth dependent pairwise amino acid substitution likelihood scores extracted from the FADHM matrices. The FADHM matrices quantify the substitution frequencies at different depths obtained by performing protein-protein structural alignments. A detailed account of the FADHM score can be found elsewhere (Farheen et al., 2017)

#### 3.2.6. The Packpred score for mutations

The Packpred score is given as,

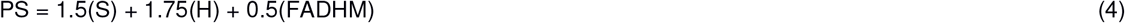

Where PS is Packpred score, S (section 3.2.5) is the residue clique score obtained from the statistical potential, H is Shannon entropy (section 3.2.3) and FADHM (section 3.2.4) is the depth based amino acid substitution likelihood score. The weights were obtained by training on T4 saturation mutagenesis data set (Supplementary Table1). The coefficients for S, H and FADHM (weights) were systematically sampled in the range 0 to 3 with a step size of 0.25. The cut off score threshold that best discriminates neutral mutations from destabilizing ones was 1.6 in the training data (see section 4.1). Mutation with a score greater than 1.6 is neutral and is destabilizing otherwise. To score a mutant, we modify the clique composition without explicitly modeling the mutant protein structure, with the mutant amino acid inheriting all the properties of the wild type residue.

Packpred is implemented as a web server at http://cospi.iiserpune.ac.in/packpred/. A standalone version is also available for download.

#### 3.2.7. Matthew’s correlation coefficient

We gauge the binary classification performance of Packpred using Matthews’s correlation coefficient (MCC) (Matthews, 1975) given as,

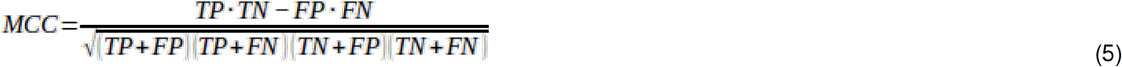

Where, TP, TN, FP and TN represents true positive, true negative, false positive and false negative predictions.

## 4. Results

### 4.1 Training and testing Packpred score

Packpred uses a linear combination of sequence position Shannon entropy, a residue clique statistical potential and a depth dependent substitution matrix (FADHM) to predict the functional effect of missense mutations. The Shannon entropy part of the score estimates the functional importance of residues based on evolutionary information. The clique statistical potential and the substitution matrix gauges the effect of the mutation on local environment/structure. The statistical potential computes the observed and expected probabilities to calculate a score for a clique. The FADHM scores are taken from substitution matrices that are derived from structural alignments of proteins. The substitution likelihood scores are calculated by categorizing a protein in three regions based on residue depths (exposed intermediate and buried). The substitution scores indicate the likelihood of a residue getting replaced by another at a given depth.

We performed a grid search in the range of 0 to 3 with a step size of 0.25 for S, H and FADHM to optimize the coefficients (weights) of each component of linear combination Packpred score. The optimization was to maximize the Matthews correlation coefficient (MCC) (see below) T4 lysozyme saturation mutagenesis data training set. The weights that gave highest MCC on the training set were 1.5, 1.75, and 0.5 for the clique statistical potential, Shannon entropy and FADHM respectively. We also obtained a cut-off threshold that distinguishes the destabilizing from the neutral ones from this training exercise. The cut off was sampled in the range 0 to 2 with a step size of 0.1. Mutations with scores greater than 1.6 are classified as neutral and scores below 1.6 are classified as destabilizing. The T4-lysozyme training set consists of 1362 (∼69%) neutral and 604 (31%) destabilizing mutations of which Packpred correctly identifies 1049(∼77%) neutral mutations and 406(∼67%) destabilizing mutations (Supplementary Table2).

The weights and threshold obtained from the training set were applied to two testing sets, CcdB saturation mutagenesis data (Supplementary Table3) and Missense3D data set (Supplementary Table4). CcdB data set has 1258 (∼80%) neutral mutations and 276 (∼20%) destabilizing while the Missense3D data set has 2134(∼52%) neutral and 1965(∼48%) disease mutations respectively. We used the PDB structures, 2LZM and 3VUB to obtain Packpred scores of T4-lysozyme and CcdB respectively. The biological unit of CcdB is a dimer and we did all the calculations using this dimeric state structure for CcdB. Packpred correctly predicts 864/1258(∼68%) neutral and 253/276(∼92%) destabilizing mutations from CcdB testing set and 1670/2134 (∼78%) neutral, 1123/1965 (∼57%) disease causing mutations from the missense3D data set.

We compared Packpred’s binary classification with several popular methods such as i-mutant2(Capriotti et al., 2005), mCSM(Pires et al., 2014b), SDM(Pandurangan et al., 2017), dynamut2(Rodrigues et al., 2020), FADHM(Farheen et al., 2017), and Missense3D (Ittisoponpisan et al., 2019) (Table 1). All the predictions were made using default parameters. Packpred was the best performing method on the T4-lysozyme training set and the Missense3D testing set with MCC values of 0.42 and 0.36 respectively. The next best method is Missense3D with MCC values of 0.40 and 0.33 for the T4 and Missense3D data sets respectively. The MCC of Packpred on the CcdB data set is 0.47 and is marginally outperformed by the best performing method, FADHM, which has an MCC of 0.48 (Table 1).

**Table 1:** Performance of some methods on T4, CcdB saturation mutagenesis and missense3D data sets. *: Values taken from FADHM paper.

| Method | MCC for T4 lysozyme saturation mutagenesis data set | MCC for CcdB saturation mutagenesis data set | MCC for Missense 3D data set |
| --- | --- | --- | --- |
| i-mutant 2.0 | 0.30* | 0.36* | 0.06 |
| mCSM | 0.22* | 0.39* | 0.05 |
| SDM2 | 0.24* | 0.33* | 0.14 |
| Dynamut2 | 0.09 | 0.15 | 0.06 |
| Missense3D | 0.40 | 0.39 | 0.33 |
| FADHM | 0.38* | **0.48\*** | 0.27 |
| Packpred | **0.42** | 0.47 | **0.36** |

### 4.2 Analysis of the predictions on the Missense3D data set

The missense3D data set has a balanced representation of ∼48% disease-associated mutations and ∼52% neutral mutations. The data set however is skewed in terms of amino-acid abundance when compared to natural abundance (Supplementary data Figure 1). For instance, Arginine has highest representation and accounts for ∼16% (664/4099) of the missense3D data set while its natural abundance is ∼5%. The next most abundant amino-acid in the Missense3D data set is glycine that accounts for ∼9% (372/4099) of the data (natural abundance is ∼7%). The most frequent mutant is also Arginine (347/4099) followed by serine (343/4099). There are 2233 mutations in the exposed environment (depth less than 5 Å), 1258 in the intermediate environment (depth between 5 and 8 Å) and 608 in the buried environment (depth greater than 8 Å).

We assessed the performance of various methods on the Missense3D data set using metrics including sensitivity, specificity, precision, recall, accuracy and f1 (Table 2). Packpred outperforms all other methods in MCC and accuracy. Missense3D has highest specificity and precision, while mCSM and i-mutant outperform all other methods in sensitivity, recall and f1-score. However, mCSM, i-mutant, SDM and dynamute predict large number of false negatives (Table 3) that affects their MCC. Hence, we compare Packpred with FADHM and Missense3D in the next sections unless otherwise stated. Packpred has less number of false positives among FADHM, Missense3D and has highest number of false negatives. The high false positive rate contributes to its lower specificity.

**Table 2:** The prediction performance of seven methods on the Missense3D data set. The best score in each assessment metric is shown in bold font.

| Metric | Packpred | FADHM | Missense3D | Dynamut2.0 | mCSM | i-mutant | SDM |
| --- | --- | --- | --- | --- | --- | --- | --- |
| MCC | **0.36** | 0.27 | 0.33 | 0.06 | 0.05 | 0.06 | 0.14 |
| Sensitivity | 0.57 | 0.39 | 0.40 | 0.84 | **0.92** | **0.92** | 0.80 |
| Specificity | 0.78 | 0.85 | **0.89** | 0.20 | 0.10 | 0.12 | 0.34 |
| Precision | 0.71 | 0.71 | **0.76** | 0.49 | 0.49 | 0.49 | 0.52 |
| Recall | 0.57 | 0.39 | 0.40 | 0.84 | **0.92** | **0.92** | 0.79 |
| Accuracy | **0.68** | 0.62 | 0.65 | 0.51 | 0.50 | 0.50 | 0.55 |
| F1 | 0.63 | 0.50 | 0.53 | 0.62 | **0.64** | **0.64** | 0.63 |

**Table 3:** Confusion matrix values for the different prediction methods. The values in bold font show the best in each category. TP, FP, TN and FN stand for True Positive, False Positive, True Negative and False Negative respectively.

| Metric | Packpred | FADHM | Missense3D | Dynamut2.0 | mCSM | i-mutant | SDM |
| --- | --- | --- | --- | --- | --- | --- | --- |
| TP | 1670 | 1816 | 1890 | 440 | 229 | 251 | 713 |
| FP | 842 | 1203 | 1177 | 312 | 158 | 164 | 420 |
| TN | 1123 | 762 | 788 | 1650 | 1804 | 1798 | 1542 |
| FN | 464 | 318 | 244 | 1685 | 1896 | 1874 | 1412 |

We analyzed the results structure wise (Supplementary Table5). Packpred correctly predicted all mutations from 264 (out of 606) structures and at least 50% mutations correctly from 507 structures. It could not correctly predict any mutation from 56 structures. In these 56 PDBs, the maximum mutations in any one protein were 4 while the average number of mutations per PDB is whole set ∼6. These 56 structures did not follow any particular discernible pattern or trait.

We stratified the missense3D data to particular depth zones to assess performance of these methods at particular depths. Packpred has 597/2233 (∼72%) correct predictions from the exposed environment, 796/1258 (∼63%) from the intermediate and 400/608 (∼66%) from the buried environment. Packpred is least accurate in predicting the effect of mutations in the intermediate environment. Interestingly, Missense3D is also the least accurate in this intermediate zone (Figure 2).

**Figure 2:**
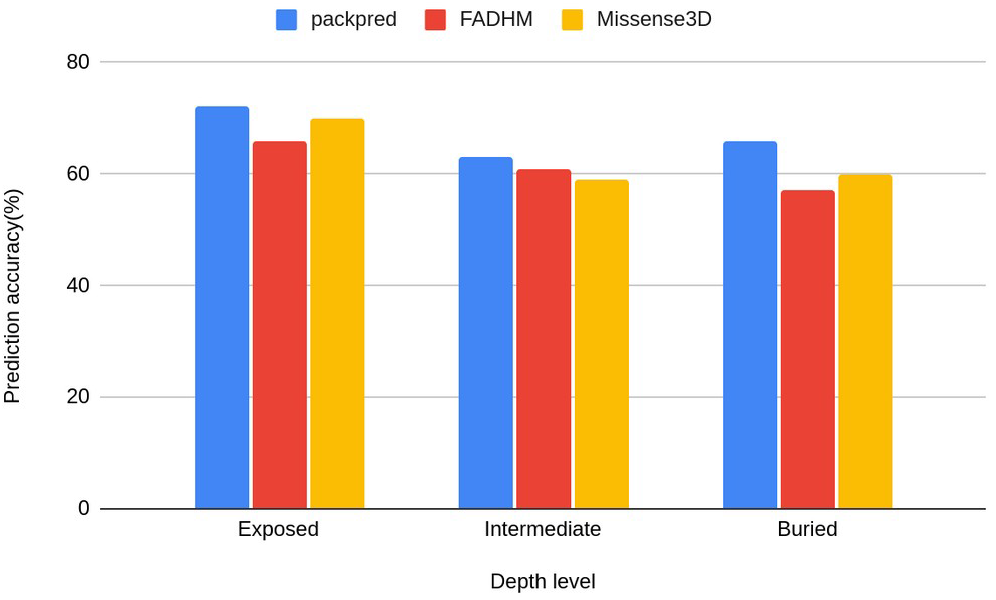
Histograms of the prediction accuracy of Packpred, FADHM and Missense3D at different depth levels (exposed to the solvent, intermediate and buried).

### 4.3 Meta predictions

Of the 4099 mutants, at least one of the seven methods we tested made an accurate prediction in 4036 cases. This motivated us to make two different meta predictions by combining the different methods.

The first meta prediction makes use of the method that performs the best for particular amino acids. We studied the wild type amino acid-wise trends of all the seven methods. For instance, amino-acids N, K, Q, R and T are best predicted by Missense3D and FADHM outperforms other methods in prediction of I and M amino acids and Packpred is best at predicting A, D, E, G, L, P, V, and Y. In fact all seven methods feature as the best method for at least one amino acid (Supplementary Table6). Interestingly, we found that Packpred has about 68% correct predictions when averaged over the 20 amino acids with a standard deviation of ∼4 while FADHM has 62% correct predictions with a standard deviation ∼7 and Missense has 64% correct predictions with ∼10 standard deviation. Dynamute2, mCSM, i-mutant2.0, and SDM have 54%, 53%, 54%, 59% correct predictions with a standard deviation of ∼ 12, 14, 14, 11 respectively. Packpred thus shows consistency in prediction across amino acid types. We then used these prediction strengths of each of the method to get a hybrid/meta prediction scheme (Supplementary Table7) that combines predictions from all of the methods and has an MCC of 0.40, easily outperforming all the individual methods.

The second meta prediction only involves Packpred, FADHM and Missense3D as these were the methods that did consistently well over all different data sets and amino acids. Here we considered the method that best predicted wild type-mutant pairs. Further, we segregated these amino acid pairs into different depth categories - exposed to solvent (depth < 5 Å), intermediate (depth between 5 to 8 Å) and buried (depth > 8 Å). Our meta prediction then chose the best performing method for a particular pair at a particular depth level. For instance, the wild type-mutant pair A→D, Packpred has the best predictions in an exposed environment, FADHM in the intermediate environment and Missense3D in the buried environment (Figure 3). In case of a tie between methods, the one with the better MCC was chosen. By thus combining the strengths of the three methods the MCC of the predictions rises to 0.51 for the Missense3D data set (Supplementary Table8). An analysis to rationalize/explain why certain methods are best for certain pairs/environments did not yield any illuminating results. It is clear however that there is some degree of complementarity in these different methods and perhaps with a more rigorous treatment of the results from the individual methods could further improve prediction accuracy.

**Figure 3:**
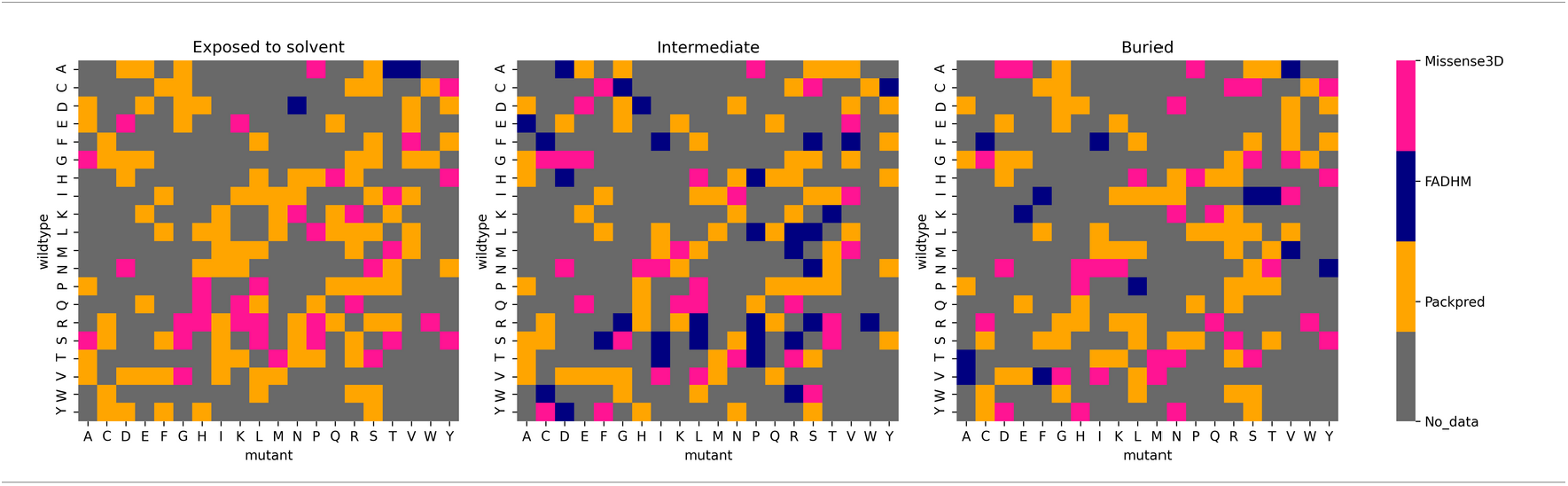
Best performing methods for each wild type-mutant amino acid pair at different depth levels.

### 4.4 Rank ordering the degree of phenotypic change by mutations

Along with the binary classification, it is also favorable for a statistical potential if it quantifies the degree of phenotypic change caused by the amino acid mutation. The degree of change is measured by the mutational sensitivity score, which categorizes each mutation into one of 4 and 8 levels in T4-lysozyme and CcdB data sets respectively. We chose to use Spearman’s rank correlation coefficient (SCC) to measure the performance of rank-ordering, as it makes no assumption on linear relationship between the scores and the phenotypical change. SCC is calculated as

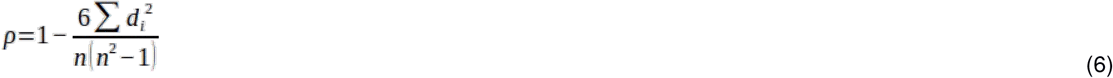

where *d* is the difference between the actual and the predicted ranks of a mutation, and *n* is the number of levels. The SCC for T4 and CcdB data sets is −0.48 and −0.54 respectively. At best, this correlation is weak and indicates that these scores could be further improved.

## 5. Discussions

In this study we have developed a method to predict what a single point mutation would do to protein function. Our method, Packpred, is constructed in a way that it is sensitive to structural changes effected by the mutation as well as any functional changes it may effect without perturbing the structures. To assess the impact of the mutation on the structure (and hence he function) of the protein, we devised a multi-body clique statistical potential. This statistical potential evaluates the strength of interaction in a local neighborhood (amino acid clique). To assess the impact of mutation, we consider the same residue neighborhood environment while replacing the wild type amino acid with the mutant. The score of the clique with the wild type residue and with the mutant are then computed. An inferior score for the mutant in comparison to the wild type would be indicative of a destabilizing mutation. The structural stability of introducing the mutant residue is also gauged by a depth dependent substitution matrix, FADHM, whose efficacy at detecting the fate of mutations we had previously benchmarked and tested. To account for functional changes that could happen even when structure is not affected on mutation, we invoke evolutionary information from a multiple sequence alignment using Shannon entropy. The more conserved the position, the more likely it is going to affect function. These different scores are taken together in a linear combination, whose coefficients were optimized using the T4-lysozyme saturation mutagenesis data set of ∼2,000 mutation.

Packpred was tested on two different data sets, another saturation mutagenesis data set (CcdB) and the Missense3D data set. Its performance on these data sets was also compared to six other method including FADHM, Missense3D, Dynamut2.0, mCSM, i-mutant2.0 and SDM. With the exception of the CcdB data set where it marginally underperforms FADHM, Packpred clearly outperformed all other methods on all data sets. Among the methods, Packpred balances well between predicting true positives and true negatives (neutral and deleterious mutations) and hence has the best MCC values. Packpred has the best accuracy and is close to the best specificity, precision and F1. It loses out to the best methods in these measures as well as on sensitivity as methods such as mCSM predict a disproportionately large number of negatives. When the performance of the different methods is compared on a (wild type) amino acid by amino acid basis, Packpred performs consistently well, with prediction accuracies never falling below 60% while maintain an average of 68%, which is easily the best among the methods tested. Qualitatively, a similar picture also emerges when the results are broken down into wild type-mutant amino acid pairs.

We also investigated whether Packpred (and other methods) preferred certain types of structures over others. No clear deduction could be made from these analyses. However, there was one trend that could be considered for further improvements – Packpred, similar to Missense3D and FADHM performed the worst in the intermediate amino acid depth environment. Mutational effects in exposed and buried (according to residue depth) environments were better predicted. Perhaps, the intermediate depth levels need to be further stratified, which in the case of Packpred would be reflected in the FADHM matrix values as well as in the clique statistical potential. Improvements could also be thought of by examining the reasons for why Packpred was unable to accurately predict the fate of 72 mutants that were all accurately called by the other six methods. We could also dissect the 23 correct predictions that Packpred made that was missed by all other method to determine the relative strength of Packpred in comparison to the other methods.

The clique statistical potential developed here has many tunable parameters such as the number of amino acids in the clique, cut off distance and definitions of what constitutes a ‘contact’ between residues, etc. Packpred could improve by investigating these aspects too and this would form an independent study in itself. Similarly, further tweaks to the FADHM matrix, as briefly discussed above, could also possibly improve overall prediction accuracy. In its current implementation, Packpred categorizes mutations as being neutral or destabilizing. When we tried to correlate the score with a discretized value of function, the correlations were around −0.5. Perhaps, with some of the improvements discussed above this correlation would also improve.

One important observations from our findings is that of the 4099 mutations, 4036 were correctly called by at least one of the methods. There exists great complementarity between the methods tested here. We were tempted to then use two simple meta prediction methods. We designated the predictions involving a particular wild type amino acid or a wild type-mutant amino acid pair to the method that best predicted this type. Such a simple minded approach gave us MCCs of 0.40 and 0.51 for the amino acid and the amino acid pair type predictions respectively, where the best predicting method, Packpred, had an MCC of 0.36 (Missense3D data set). It is conceivable that a different method of combining the results from these different method could vastly increase accuracy of predicting the functional fate of single amino acid changes.

## 6. Acknowledgments

M.S.M. would like to acknowledge the DBT-Wellcome India alliance for a senior fellowship. We would like to thank Dr. Chandra Verma and Prof. Raghavan Varadarajan and members of the COSPI lab for discussions and constructive criticisms. We thank Gulzar Singh and Parichit Sharma for their contribution to the Packpred web server. We also thank Swastik Mishra and Neelesh Soni for their technical assistance towards the Packpred software.

## References

Adkar, B. V., Tripathi, A., Sahoo, A., Bajaj, K., Goswami, D., Chakrabarti, P., et al. (2012). Protein model discrimination using mutational sensitivity derived from deep sequencing. Structure 20, 371–381. doi:10.1016/j.str.2011.11.021.

Adzhubei, I. A., Schmidt, S., Peshkin, L., Ramensky, V. E., Gerasimova, A., Bork, P., et al. (2010). A method and server for predicting damaging missense mutations. Nat. Methods 7, 248–249. doi:10.1038/nmeth0410-248.

Altschul, S. F., Madden, T. L., Schäffer, A. A., Zhang, J., Zhang, Z., Miller, W., et al. (1997). Gapped BLAST and PSI-BLAST: A new generation of protein database search programs. Nucleic Acids Res. 25, 3389–3402. doi:10.1093/nar/25.17.3389.

Altshuler, D. M., Durbin, R. M., Abecasis, G. R., Bentley, D. R., Chakravarti, A., Clark, A. G., et al. (2012). An integrated map of genetic variation from 1,092 human genomes. Nature 491, 56– 65. doi:10.1038/nature11632.

Bateman, A., Martin, M. J., O’Donovan, C., Magrane, M., Alpi, E., Antunes, R., et al. (2017). UniProt: The universal protein knowledgebase. Nucleic Acids Res. 45, D158–D169. Doi:10.1093/nar/gkw1099.

Berman, H. M., Westbrook, J., Feng, Z., Gilliland, G., Bhat, T. N., Weissig, H., et al. (2000). The Protein Data Bank. Nucleic Acids Res. 28, 235–242. doi:10.1093/nar/28.1.235.

Capriotti, E., Fariselli, P., and Casadio, R. (2005). I-Mutant2.0: Predicting stability changes upon mutation from the protein sequence or structure. Nucleic Acids Res. 33. doi:10.1093/nar/gki375.

Chakravarty, S., and Varadarajan, R. (1999). Residue depth: A novel parameter for the analysis of protein structure and stability. Structure 7, 723–732. doi:10.1016/S0969-2126(99)80097-5.

Craig Venter, J., Adams, M. D., Myers, E. W., Li, P. W., Mural, R. J., Sutton, G. G., et al. (2001). The sequence of the human genome. Science (80-.). 291, 1304–1351. doi:10.1126/science.1058040.

Dehouck, Y., Grosfils, A., Folch, B., Gilis, D., Bogaerts, P., and Rooman, M. (2009). Fast and accurate predictions of protein stability changes upon mutations using statistical potentials and neural networks: PoPMuSiC-2.0. Bioinformatics 25, 2537–2543. doi:10.1093/bioinformatics/btp445.

Farheen, N., Sen, N., Nair, S., Tan, K. P., and Madhusudhan, M. S. (2017). Depth dependent amino acid substitution matrices and their use in predicting deleterious mutations. Prog. Biophys. Mol. Biol. 128, 14–23. doi:10.1016/j.pbiomolbio.2017.02.004.

Finn, R. D., Bateman, A., Clements, J., Coggill, P., Eberhardt, R. Y., Eddy, S. R., et al. (2014). Pfam: The protein families database. Nucleic Acids Res. 42, D222. Doi:10.1093/nar/gkt1223.

Frazer, K. A., Ballinger, D. G., Cox, D. R., Hinds, D. A., Stuve, L. L., Gibbs, R. A., et al. (2007). A second generation human haplotype map of over 3.1 million SNPs. Nature 449, 851–861. doi:10.1038/nature06258.

Ittisoponpisan, S., Islam, S. A., Khanna, T., Alhuzimi, E., David, A., and Sternberg, M. J. E. (2019). Can Predicted Protein 3D Structures Provide Reliable Insights into whether Missense Variants Are Disease Associated? J. Mol. Biol. 431, 2197–2212. doi:10.1016/j.jmb.2019.04.009.

Karczewski, K. J., Weisburd, B., Thomas, B., Solomonson, M., Ruderfer, D. M., Kavanagh, D., et al. (2017). The ExAC browser: Displaying reference data information from over 60 000 exomes. Nucleic Acids Res. 45, D840–D845. Doi:10.1093/nar/gkw971.

Landrum, M. J., Lee, J. M., Riley, G. R., Jang, W., Rubinstein, W. S., Church, D. M., et al. (2014). ClinVar: Public archive of relationships among sequence variation and human phenotype. Nucleic Acids Res. 42, D980. Doi:10.1093/nar/gkt1113.

Loris, R., Dao-Thi, M. H., Bahassi, E. M., Van Melderen, L., Poortmans, F., Liddington, R., et al. (1999). Crystal structure of CcdB, a topoisomerase poison from E. coli. J. Mol. Biol. 285, 1667– 1677. doi:10.1006/jmbi.1998.2395.

Masso, M., and Vaisman, I. I. (2014). AUTO-MUTE 2.0: A portable framework with enhanced capabilities for predicting protein functional consequences upon mutation. Adv. Bioinformatics 2014. doi:10.1155/2014/278385.

Ng, P. C., and Henikoff, S. (2003). SIFT: Predicting amino acid changes that affect protein function. Nucleic Acids Res. 31, 3812–3814. doi:10.1093/nar/gkg509.

Pandurangan, A. P., Ochoa-Montaño, B., Ascher, D. B., and Blundell, T. L. (2017). SDM: A server for predicting effects of mutations on protein stability. Nucleic Acids Res. 45, W229– W235. Doi:10.1093/nar/gkx439.

Pires, D. E. V., Ascher, D. B., and Blundell, T. L. (2014a). DUET: A server for predicting effects of mutations on protein stability using an integrated computational approach. Nucleic Acids Res. 42. doi:10.1093/nar/gku411.

Pires, D. E. V., Ascher, D. B., and Blundell, T. L. (2014b). MCSM: Predicting the effects of mutations in proteins using graph-based signatures. Bioinformatics 30, 335–342. doi:10.1093/bioinformatics/btt691.

Rennell, D., Bouvier, S. E., Hardy, L. W., and Poteete, A. R. (1991). Systematic mutation of bacteriophage T4 lysozyme. J. Mol. Biol. 222. doi:10.1016/0022-2836(91)90738-R.

Roach, J. C., Glusman, G., Smit, A. F. A., Huff, C. D., Hubley, R., Shannon, P. T., et al. (2010). Analysis of genetic inheritance in a family quartet by whole-genome sequencing. Science (80-.). 328, 636–639. doi:10.1126/science.1186802.

Rodrigues, C. H. M., Pires, D. E. V., and Ascher, D. B. (2020). DynaMut2: Assessing changes in stability and flexibility upon single and multiple point missense mutations. Protein Sci., pro.3942. Doi:10.1002/pro.3942.

Šali, A., and Blundell, T. L. (1993). Comparative protein modelling by satisfaction of spatial restraints. J. Mol. Biol. 234, 779–815. doi:10.1006/jmbi.1993.1626.

Sippl, M. J. (1990). Calculation of conformational ensembles from potentials of mena force. An approach to the knowledge-based prediction of local structures in globular proteins. J. Mol. Biol. 213, 859–883. doi:10.1016/S0022-2836(05)80269-4.

Smith, H. O., Annau, T. M., and Chandrasegaran, S. (1990). Finding sequence motifs in groups of functionally related proteins. Proc. Natl. Acad. Sci. U. S. A. 87, 826–830. doi:10.1073/pnas.87.2.826.

Stranger, B. E., Stahl, E. A., and Raj, T. (2011). Progress and promise of genome-wide association studies for human complex trait genetics. Genetics 187, 367–383. doi:10.1534/genetics.110.120907.

Tan, K. P., Nguyen, T. B., Patel, S., Varadarajan, R., and Madhusudhan, M. S. (2013). Depth: a web server to compute depth, cavity sizes, detect potential small-molecule ligand-binding cavities and predict the pKa of ionizable residues in proteins. Nucleic Acids Res. 41. doi:10.1093/nar/gkt503.

Tan, K. P., Varadarajan, R., and Madhusudhan, M. S. (2011). DEPTH: A web server to compute depth and predict small-molecule binding cavities in proteins. Nucleic Acids Res. 39, W242– W248. Doi:10.1093/nar/gkr356.

Weaver, L. H., and Matthews, B. W. (1987). Structure of bacteriophage T4 lysozyme refined at 1.7 Å resolution. J. Mol. Biol. 193, 189–199. doi:10.1016/0022-2836(87)90636-X.

Worth, C. L., Preissner, R., and Blundell, T. L. (2011). SDM – A server for predicting effects of mutations on protein stability and malfunction. Nucleic Acids Res. 39, W215. Doi:10.1093/nar/gkr363.

Yates, C. M., Filippis, I., Kelley, L. A., and Sternberg, M. J. E. (2014). SuSPect: Enhanced prediction of single amino acid variant (SAV) phenotype using network features. J. Mol. Biol. 426, 2692–2701. doi:10.1016/j.jmb.2014.04.026.

Zhang, Z., Miteva, M. A., Wang, L., and Alexov, E. (2012). Analyzing effects of naturally occurring missense mutations. Comput. Math. Methods Med. 2012, 15. doi:10.1155/2012/805827.

Altschul, S.F., Madden, T.L., Schäffer, A.A., Zhang, J., Zhang, Z., Miller, W. & Lipman, D.J. (1997). Gapped BLAST and PSI-BLAST: a new generation of protein database search programs. Nucleic Acids Res. 25:3389–3402.

