## Supplementary data Figure 1 for "Packpred: Predicting the functional effect of missense mutations"

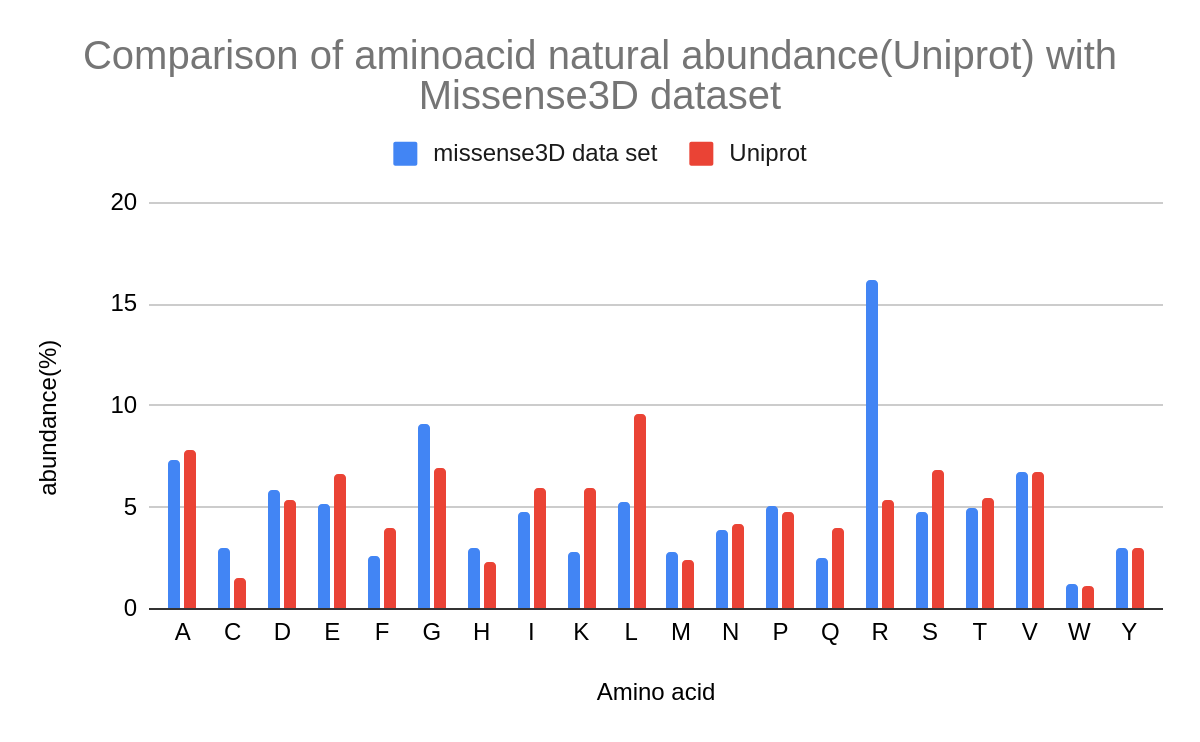

Supplementary Figure1: Abundance of amino-acids in Missense3D dataset in comparison to their natural abundance (Taken from UniprotKB 8.0)
